# Early diverging insect pathogenic fungi of the order *Entomophthorales* possess diverse and unique subtilisin-like serine proteases

**DOI:** 10.1101/247858

**Authors:** Jonathan A. Arnesen, Joanna Malagocka, Andrii Gryganskyi, Igor V. Grigoriev, Kerstin Voigt, Jason E. Stajich, Henrik H. De Fine Licht

**Affiliations:** Section for Organismal Biology, Department of Plant and Environmental Sciences, University of Copenhagen, Thorvaldsenvej 40, 1871 Frederiksberg, Denmark.; Department of Biology, Duke University, Durham, North Carolina, USA.; US Department of Energy Joint Genome Institute, Walnut Creek, California, USA.; Jena Microbial Resource Collection (JMRC), Leibniz Institute for Natural Product Research and Infection Biology - Hans Knoell Institute, Jena, Germany.; Institute of Microbiology, Friedrich Schiller University, Jena, Germany.; Department of Plant Pathology and Microbiology, University of California, Riverside, California, USA.

**Keywords:** Subtilase, insect-pathogen, early-diverging fungi, proteases

## Abstract

Insect-pathogenic fungi use subtilisin-like serine proteases (SLSPs) to degrade chitin-associated proteins in the insect procuticle. Most insect-pathogenic fungi in the order Hypocreales (Ascomycota) are generalist species with a broad host-range, and most species possess a high number of SLSPs. The other major clade of insect-pathogenic fungi is part of the subphylum Entomophthoromycotina (Zoopagomycota, formerly Zygomycota) which consists of high host-specificity insect-pathogenic fungi that naturally only infect a single or very few host species. The extent to which insect-pathogenic fungi in the order Entomophthorales rely on SLSPs is unknown. Here we take advantage of recently available transcriptomic and genomic datasets from four genera within Entomophthoromycotina: the saprobic or opportunistic pathogens *Basidiobolus meristosporus, Conidiobolus coronatus, C. thromboides, C. incongruus,* and the host-specific insect pathogens *Entomphthora muscae* and *Pandora formicae,* specific pathogens of house flies (*Muscae domestica*) and wood ants (*Formica polyctena*), respectively. We use phylogenetics and protein domain analysis to show that the obligate biotrophic fungi *E. muscae, P. formicae* and the saprobic human pathogen *C. incongruus* all contain “classical” fungal SLSPs and a unique group of SLSPs that loosely resembles bacillopeptidase F-like SLSPs. This novel group of SLSPs is found in the genomes of obligate insect pathogens and a generalist saprobic opportunistic pathogen why they are unlikely to be responsible for the host specificity of Entomophthorales. However, this class represent a unique group of SLSPs so far only observed among Bacteria, Oomycetes and early diverging fungi such as Cryptomycota, Microsporidia, and Entomophthoromycotina and missing in the sister fungal lineages of Kickxellomycotina or the fungal phyla Mucoromyocta, Ascomycota and Basidiomycota fungi suggesting interesting gene loss patterns.

## 1. Introduction

Insect pathogenic fungi use a broad array of enzymes to penetrate the host cuticle and gain entry to the soft tissues inside (Charnley, 2003; St. Leger et al., 1986b). In many cases, serine proteases are among the first enzymes to be secreted in the early stages of infection in order to cleave and open up chitin-associated proteins in the procuticle (St. Leger et al., 1986a; Vega et al., 2012), which later is followed by extensive lipase and chitinase enzymatic secretions (Charnley, 2003). In particular, subtilisin-like serine proteases (SLSPs) have been considered important virulence factors in pathogenic fungi (Muszewska et al., 2011). The first SLSPs from insect pathogenic fungi were identified in *Metarhizium anisopliae* (ARSEF2575), which secretes SLSPs as some of the key proteases during fungal growth on insect cuticle (Charnley, 2003; St. Leger et al., 1986a). Comparative genomic approaches have identified significant expansions of the SLSP gene family in the genus *Metarhizium* (Bagga et al., 2004; Hu et al., 2014), the insect pathogenic fungus *Beauveria bassiana* (Xiao et al., 2012), two nematode-trapping fungi *Monacrosporium haptotylum* and *Arthrobotrys oligospora* that are able to penetrate the chitinaceous cell wall of soil nematodes (Meerupvati et al. 2013), and dermatophytic fungi such as *Arthroderma benhamiae* and *Trichophyton verrucosum* that can cause nail and skin infections in humans and animals (Burmester et al., 2011; Desjardins et al., 2011; Martinez et al., 2012; Sharpton et al., 2009). Fungi that are able to utilize chitin-rich substrates, including many insect pathogenic fungi, thus appear to often be associated with a diversified and expanded set of SLSPs.

Although SLSPs are expanded among insect pathogenic fungi, this group of proteases are ubiquitous among eukaryotic organisms. Most SLSPs are secreted externally or localized to vacuoles, and especially in saprobic and symbiotic fungi SLSPs constitute an important component of the secretome (Li et al., 2017). According to the MEROPS peptidase classification, the S8 family of SLSPs together with the S53 family of serine-carboxyl proteases make up the SB clan of subtilases (Rawlings et al., 2016). The S8 family of SLSPs is characterized by an Asp-His-Ser catalytic triad (DHS triad), which forms the active site and is further divided into two subfamilies S8A and S8B. Subfamily S8A contains most S8 representatives, including the well-known Proteinase K enzyme that is widely used in laboratories as a broad-spectrum protease. The S8B SLSPs are kexins and furins which cleave peptides and protohormones (Jalving et al., 2000; Muszewska et al., 2017, 2011). Based on characteristic protein domain architectures and protein motifs surrounding the active site residues, the large S8A subfamily of SLSPs is further divided into several groups such as proteinase-K and pyrolysin. Besides these two major groups of proteinase K-like and pyrolisin subfamilies, six new groups of subtilase genes designated *new 1* to *new 6* have recently been found (Li et al., 2017; Muszewska et al., 2011). The analysis of fungal genome data from a wide taxonomic range has shown that the size of the proteinase K gene family has expanded independently in fungi pathogenic to invertebrates (Hypocreales) and vertebrates (Onygenales) (Muszewska et al., 2017; Sharpton et al., 2009). Closely related systemic human-pathogenic fungi, however, do not show the same expansions and related pathogens and non-pathogens can show the same expansions (Muszewska et al., 2011; Whiston and Taylor, 2016). This suggests that the number of SLSPs that a fungus possess is not directly related to pathogenicity, but instead is associated with the use of dead or alive animal tissue as growth substrate (Li et al., 2017; Muszewska et al., 2011).

Most anamorphic insect-pathogenic fungi in the order Hypocreales (Ascomycota) are generalist species with a broad host-range capable of infecting most major orders of insects (e.g. *M. robertsii* and *B. bassiana*) or specific to larger phylogenetic groups (e.g. the locust-specific *M. acridum* or the coleopteran pathogen *B. brongniartii*). The above inferences of fungal SLSP evolution rely almost exclusively on insights from Ascomycota, and consequently has strong sampling bias towards generalist insect-pathogenic fungi. In contrast, the other major clade of insect-pathogenic fungi in the subphylum Entomophthoromycotina (Zoopagomycota, formerly Zygomycota) consists almost exclusively of insect-pathogens and many are extremely host-specific, naturally only infecting a single or very few host species (Spatafora et al., 2016). The dearth of genomic data for Entomophthoromycotina has previously precluded their inclusion in comparative genomic analyses (De Fine Licht et al., 2016; Gryganskyi and Muszewska, 2014). Here we take advantage of recently available transcriptomic and genomic datasets from four genera within Entomophthoromycotina: the saprobic *Basidiobolus meristosporus,* the saprobic and opportunistic pathogens, *Conidiobolus coronatus, C. thromboides, C. incongruus*, and the host-specific insect pathogens *Entomphthora muscae* and *Pandora formicae,* specific pathogens of house flies (*Muscae domestica*) and wood ants (*Formica polyctena*), respectively. We use phylogenetics and protein domain analysis to show that the obligate biotrophic fungi *E. muscae, P. formicae* and the saprobic human pathogen *C. incongruus* in addition to more “classical” fungal SLSPs, harbor a unique group of SLSPs that loosely resembles bacillopeptidase F-like SLSPs.

## 2. Materials and Methods

### Sequence database searches for subtilisin-like serine proteases

We identified putative subtilisin-like serine proteases (SLSPs) from six fungi in the order Entomophthorales: *Entomophthora muscae, Pandora formicae, Basidiobolus meristosporus, Conidiobolus coronatus, C. incongruus* and *C. thromboides*. First, Pfam protein family domains were identified in the *de-novo* assembled transcriptomes of *E. muscae* KVL-14-117 (De Fine Licht et al., 2017), and *P. formicae* (Małagocka et al., 2015), using profile Hidden Markov Models with hmmscan searches (e-value < 1e-10) against the PFAM-A database ver. 31.0 using HMMER ver. 3 (Eddy, 1998; Finn et al., 2016). All sequences in the transcriptome datasets containing the S8 subtilisin-like protease domain (PF00082) were identified and included in further analyses. Second, all sequences that contain the PF00082 domain were obtained from the genomes of *B. meristosporus* CBS 931.73 (Mondo et al., 2017), *C. coronatus* NRRL28638 (Chang et al., 2015), and *C. thromboides* FSU 785 from the US Department of Energy Joint Genome Institute *MycoCosm* genome portal (http://jgi.doe.gov/-fungi). Third, predicted coding regions in the genome sequence of *C. incongruus* CDC-B7586 (Chibucos et al., 2016), were searched for the presence of the S8 subtilisin-like protease domain (PF00082) as described above.

Sequences encoding an incomplete Asp-His-Ser catalytic triad (DHS triad) characteristic of S8 family proteases were regarded as potential pseudogenes and excluded from further analysis as they. A preliminary protein alignment made with ClustalW and construction of a Neighbour-Joining tree revealed a highly divergent group of *P. formicae* SLSP-sequences that had significant homology with insect proteases (blastp, e-value < 1e-6, ncbi-nr protein database, accessed June 2017). These putative insect-sequences likely originate from the ant host, *Formica polyctena,* and represents host contamination that were not filtered out from the dual-RNAseq reads used to construct the *P. formicae* transcriptome (Małagocka et al., 2015). These divergent sequences were therefore removed and excluded from further analysis.

### Protein domain architecture and homology modelling

The domain architectures of all putative SLSPs identified within Entomophthoromycotina were predicted using Pfam domain annotation. The presence of putative signal peptides for extracellular secretion were predicted using SignalP (Petersen et al., 2011). Homology-based protein models of the 3D structure were constructed with Swiss-model (Biasini et al., 2014) and visualized using PyMol (The PyMOL Molecular Graphics System, Version 1.8 Schrödinger, LLC).

### Protease sequence clustering

A Markov Clustering Algorithm (MCL) was used to identify clusters of similar proteins among putative SLSPs identified within Entomophthorales. Clustering using MCL is based on a graph constructed by an all-vs-all-BLAST of SLSPs (BLASTP, e-value < 1e^−10^). The Tribe-MCL protocol (Enright et al., 2002) as implemented in the Spectral Clustering of Protein Sequences (SCPS) program (Nepusz et al., 2010) was used with *inflation* = 2.0. The *inflation* parameter is typically set between 1.2 – 5.0 (Nepusz et al., 2010), and controls the “tightness” of the sequence matrix, with lower values leading to fewer clusters and higher values to more sequence clusters. To putatively assign protease function to the newly identified Entomophthorales sequence clusters the Tribe-MCL protocol was used to identify clusters of similar proteins between the putative SLSPs identified within Entomophthorales and 20,806 protease sequences belonging to the peptidase subfamily S8A obtained from the MEROPS database, accessed November 2017 (Rawlings et al., 2016). Investigation of the MEROPS protease sequences that clustered together with the identified Entomophthorales sequence clusters allowed putative protease holotype information to be assigned to the identified clusters.

### Phylogenetic analysis

All identified putative Entomophthorales SLSP coding nucleotide sequences were aligned in frame to preserve codon structure using MAFFT (Larkin et al., 2007). Unreliable codon-columns with a Guidance2 score below 0.90 in the multiple sequence alignment were removed (Penn et al., 2010). The best model for phylogenetic analysis was selected by running PhyML with *GTR* as substitution model and with-or-without Gamma parameter and a proportion of invariable sites (Guindon and Gascuel, 2003). The optimal substitution model based on the Bayesian Information Criterion (BIC) score (*GTR+G*) was used in maximum likelihood analysis calculation using RaxML with 10,000 bootstrap runs (Stamatakis, 2014).

To identify branches that potentially contain signatures of positive selection among SLSP sequences we used maximum likelihood estimates of the dN/dS ratio (ω) for each site (codon) along protein sequences. A specific lineage (branch) was tested independently for positive selection (ω > 1) on individual sites by applying a neutral model that allows ω to vary between 0 – 1 and a selection model that also incorporates sites with ω > 1 using the software *codeml* implemented in PAML 4.4 (Yang, 2007). Statistical significance was determined with a likelihood ratio test of these two models for the tested lineage.

## 3. Results

We identified 154 SLSP sequences from six fungi in the subphylum Entomophthoromyctina: *E. muscae* (n = 22), *P. formicae* (n = 6), *B. meristosporus* (n = 60), *C. thromboides* (n = 18), *C. coronatus* (n = 36), and *C. incongruus* (n = 12). Close inspection of the active site residues revealed two *C. incongruus* sequences (Ci7229 and Ci12055), which contained the active site DHS residues in the motifs <u>Asp</u>-Asp-Gly, <u>His</u>-Gly-Thr-Arg, and Gly-Thr-<u>Ser</u>-Ala/Val-Ala/Ser-Pro characteristic of the S8B subfamily of S8 proteases. These two sequences also contained the P domain (PF01483) indicating that they are S8B kexin proteases. All other 152 identified Entomophthoromyctina S8 protease sequences contained active site residues closely resembling the motifs Asp-Thr/Ser-Gly, His-Gly-Thr-His, and Gly-Thr-Ser-Met-Ala-Xaa-Pro characteristic of the S8A subfamily. Cluster analysis of these 152 S8A-protease sequences identified three groups of proteins that were designated as group A, B and C, with 130, 11 and 11 sequences in each cluster, respectively (Fig. 1). These groups do not change when the Tribe-MCL *inflation* parameters varied within a range of 1.5 – 6.0 suggesting that these three groups were clearly distinct. Phylogenetic analysis of the identified 152 S8A SLSPs using maximum likelihood methods (Stamatakis, 2014) also recovered the same three distinct lineages with strong bootstrap support (Fig. 1). Evidence of positive selection acting on specific enzyme residues on the branches leading to these clusters were not detected with branch-site tests (2ΔlnL > 1.53, P > 0.217).

**Figure 1.**
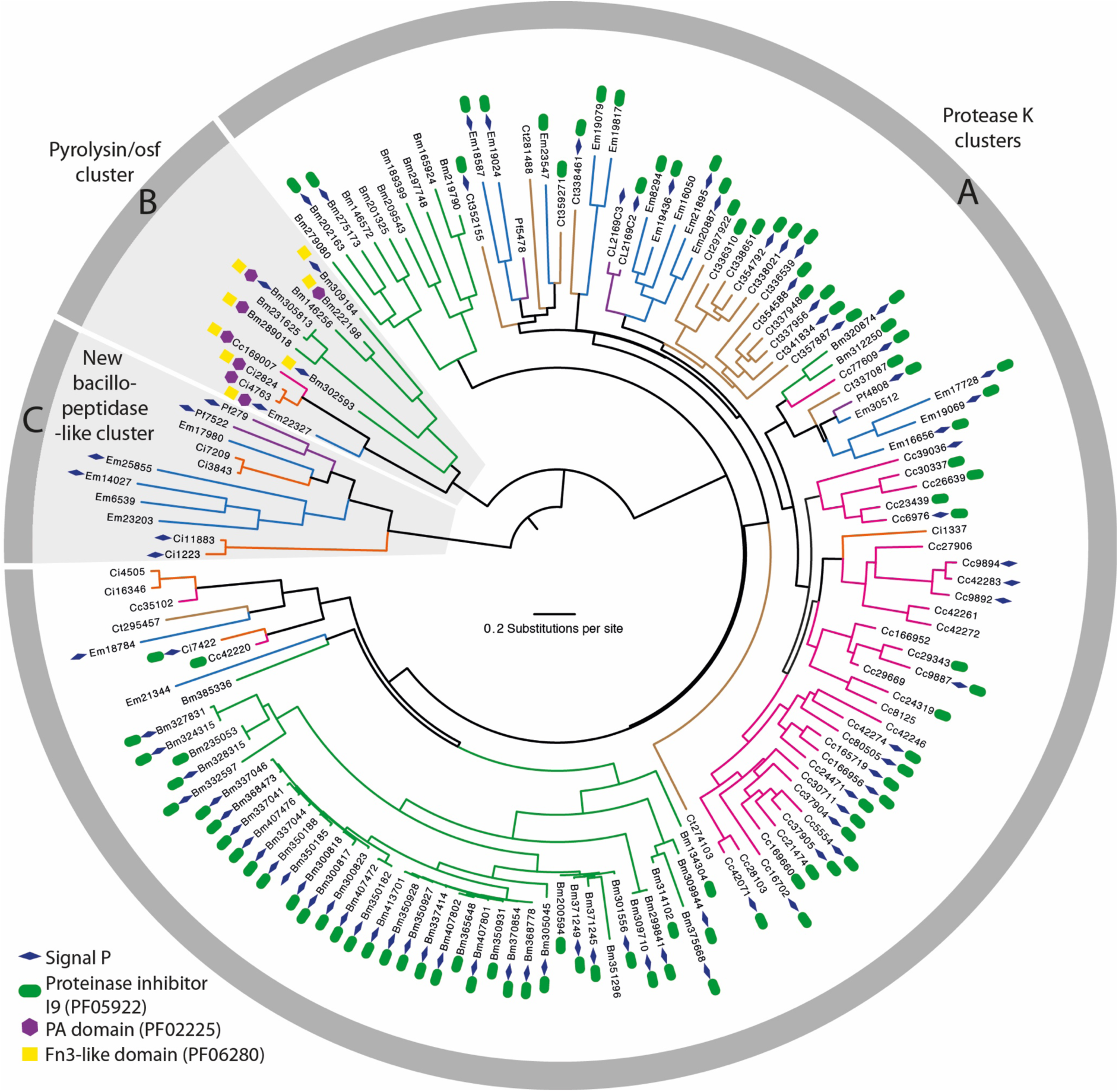
Maximum likelihood phylogeny calculated with RAxML and based on a 2379 bp alignment of 152 subtilisin-like serine protease codon nucleotide sequences from Entomophthoromycotina that contain the peptidase S8/S53-subtilisin (PF00082) domain. Branches are coloured for eachs species as *Entomophthora muscae* (Blue), *Pandora formicae* (Purple), *Conidiobolus coronatus* (Pink), *C. thromboides* (brown), *C. incongruus* (orange), and *B. meristopolus* (Green). For each SLSP, the accession number and protein domains additional to PF00082 are shown. The three identified clusters: Protease K cluster (A), Pyrolysin/osf protease cluster (B), and the new bacillopeptidase-like Entomophthorales cluster (C), are marked in the grey circle surrounding the tree and with grey background for cluster B and C.

To further characterize the three SLSP groups, the protein domain architecture of each of the 152 protease sequences were analyzed. The presence of a proteinase-associated (PA, pfam:PF02225) domain was only found in Group B, strongly suggested that this cluster with 11 members is comprised of pyrolisin and osf proteases (Muszewska et al., 2011). An additional Tribe-MCL cluster analysis (inflation = 1.2) of the 152 Entomophthoromycotina and all S8A proteases in the MEROPS database clustered these 11 Entomophthoromycotina sequences into a group of 779 MEROPS proteases. This group of proteases contained 42 members of the fungal S08.139 (PoSl-(*Pleurotus ostreatus*)-type peptidase) holotype (Supplementary data). The protein domain architecture of group A contained 130 Entomopthoromycotina SLSPs with secretory signal and a peptidase inhibitor (Inhibitor_I9, pfam: PF05922) domain. None of the 11 members of group C contained a peptidase inhibitor domain (Fig. 1). The 130 Entomophthoromycotina Group B SLSPs appear to belong to the common Proteinase K group of S8A proteases (Muszewska et al., 2011) based on conservation of the active site residues (Fig. 2) and cluster membership of the MEROPS Tribe-MCL analysis. The 130 Entomophthoromycotina SLSPs clustered with 3,046 MEROPS proteins of well-known fungal entomopathogenic protease holotypes (Supplementary data), including cuticle-degrading peptidase of nematode-trapping fungi (S08.120), cuticle-degrading peptidase of insect-pathogenic fungi in the genus *Metarhizium* (S08.056), and subtilisin-like peptidase 3 (*Microsporum-*type; S08.115).

**Figure 2.**
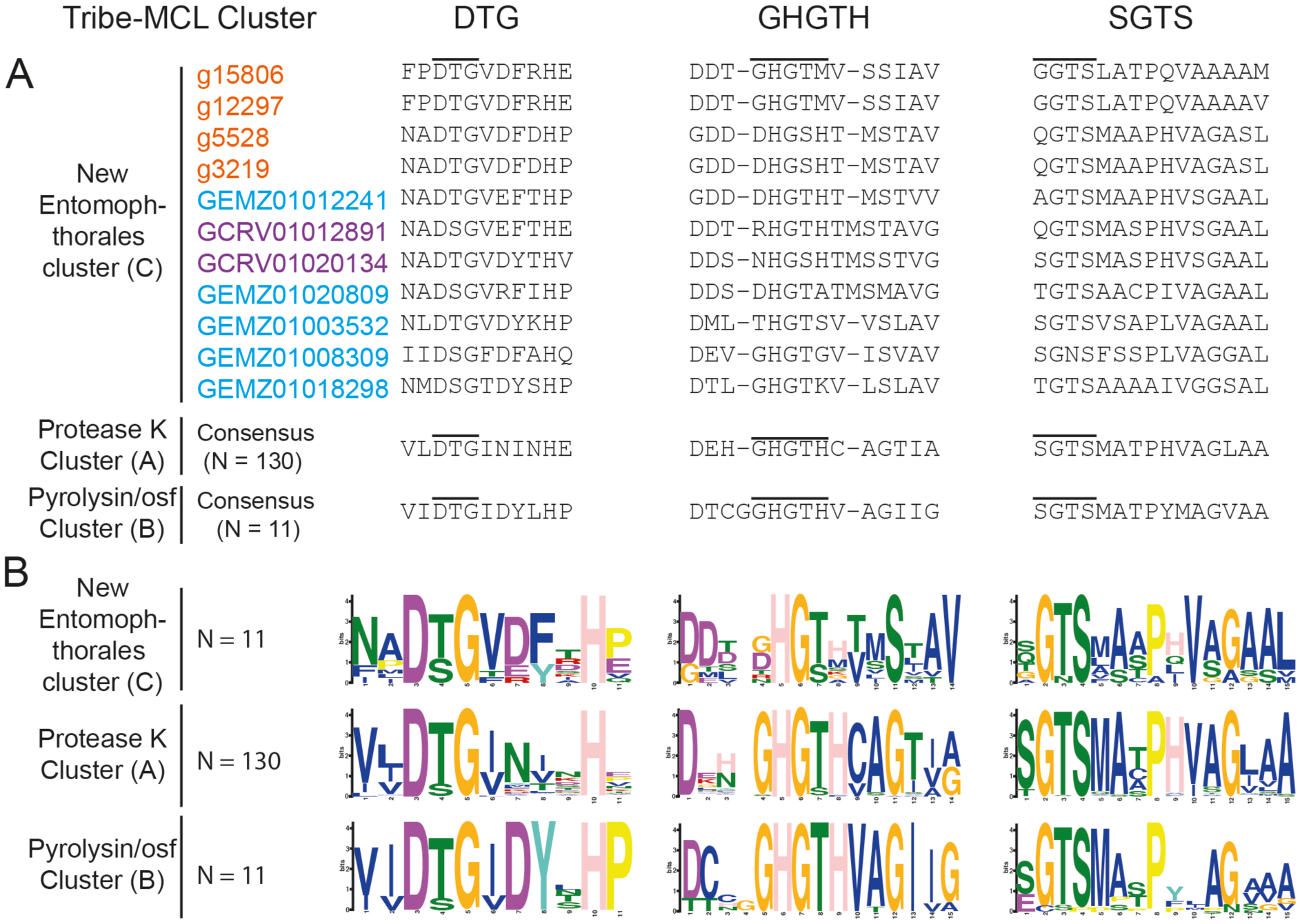
Active site and domain co-occurrence variability of the three Tribe-MCL clusters identified among 152 Entomophthoromycotina subtilisin-like serine proteases. The columns DTG, GHGTH, and SGTS represents the closest amino acid sequence for each of the amino acids from the DHS catalytic triad. **A.** Amino acid alignment of the active site residues for the three identified groups (A-C) of SLSPs within Entomophthoromycotina. Accession codes are color coded as: Orange – *C. incongruus,* Blue – *E. muscae,* and Purple – *P. formicae.* **B.** Sequence motifs of the active site residues for each group.

The MEROPS Tribe-MCL analysis identified a third group (C) of 11 Entomophthoromycotina SLSPs which were similar to 402 MEROPS proteins. Of these members 386 were classified as unassigned subfamily S8A peptidases (S08.UPA) and the remaining 15 assigned to the bacillopeptidase F holotype (S08.017; Supplementary data). All 402 MEROPS proteases in this group originated from either Bacteria or Oomycetes, except for two proteases from the Fungi *Rozella allomycis* (Cryptomycota) and *Mitosporidium daphnia* (Microsporidia), respectively (Fig. 3, Supplementary data). The group C Entomophthorales SLSPs clustered as sister to these two Cryptomycota and Microsporidia proteins with strong support (ML bootstrap value = 99) in the phylogenetic analysis of the protein sequences (Fig. 3), in concordance with this group of SLSPs being an outlier from all other previously known fungal S8A SLSPs.

**Figure 3.**
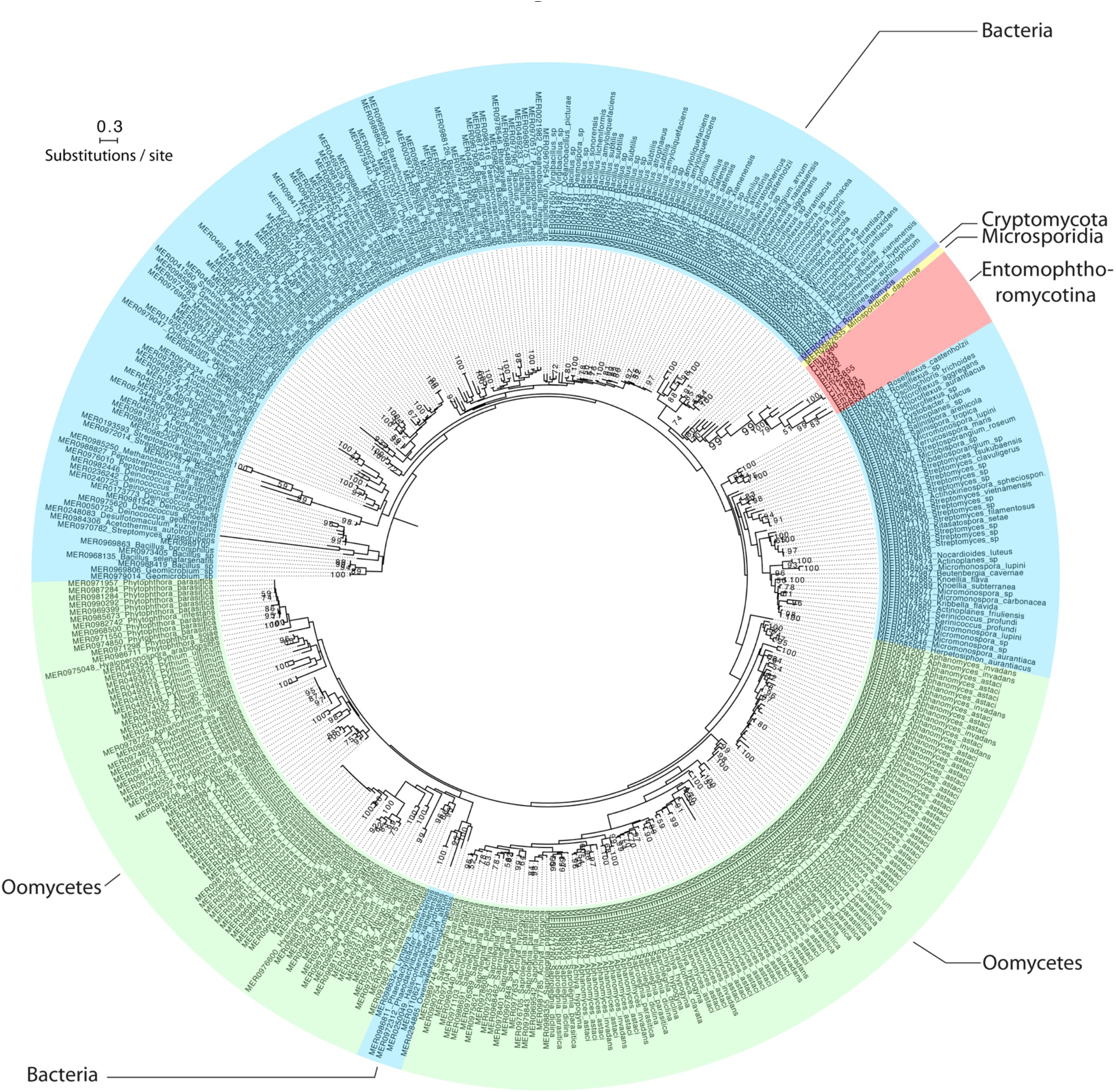
Mid-point rooted maximum likelihood phylogeny calculated with RAxML and based on a (479 amino acid) alignment of 413 protein subtilisin-like serine protease sequences, which belonged to group C in the Tribe-MCL analysis (see text for details). Bootstrap values >50 from 1000 iterations are shown.

## 4. Discussion

Subtilisin-like serine proteases (SLSPs) have many roles in fungal biology and are known to be involved in host–pathogen interactions. Independent expansion of copy number and diversification of SLSPs is widespread among animal pathogenic Dikarya (Ascomycota and Basidiomycota) (Li et al., 2017). The repeated expansion of SLSPs among the generalist insect-pathogenic hypocrealean fungi has been interpreted as an adaptation to enable infection of insect hosts (Muszewska et al., 2011), whereas comparatively little is known about the evolution and diversification of SLSPs among the early diverging fungal clades. To understand the evolution of SLSPs among the vertebrate and arthropod pathogenic fungi in the subphylum Entomophthoromycotina, we searched available genomic and transcriptomic sequence data to identify all entomophthoralean genes with SLSP domains. We found 154 entomophthoromycotan SLSPs, of which two copies were classified as S8B kexin SLSPs.

The remaining 152 S8A SLSPs were clustered by sequence similarity and compared by phylogenetic analysis to show that the majority of the SLSPs (n = 130) are similar to and cluster together with “classical” proteinase-K-like fungal S8A SLSPs (Figure 1).

We did identify 11 SLSPs that cluster together with 402 un-annotated or Bacillopeptidase F-like SLSPs from bacteria and Oomycetes (Figure 3). These obervations remained consistent even when exploring variation in the inflation parameter, which controls the “tightness” in the cluster analysis. The entomophthoralean and Oomycete S8A SLSPs form separate clades within this cluster of primarily bacterial proteases indicating that the Entomophthorales and Oomycete SLSPs evolved independently (Figure 3). Apart from the entomophthoralean sequences, only two fungal protease sequences were found within this group from the Cryptomycota *R. allomycis* and microsporidium *M. daphnia.* A statistical test for a significant expansion of SLSP copy number among the insect-pathogenic Entomophthorales was not explicitly performed in the present analysis due to uncertainty of total gene numbers from transcriptomic data sets of the specialist insect-pathogens *E. muscae* and *P. formicae.* In the sampled transcriptomes, the number of transcripts is likely larger than the genome gene count due to splice variants, post-transcriptional modifications, and allelic variants assembling into multiple transcripts per gene. In addition, the assembled transcripts only reflect actively transcribed genes expressed in the sampled conditions and time points, and may underestimate the actual number of genes. These confounding factors impact the estimated number of genes and make quantitative comparative analyses of gene family size between transcriptomes unreliable. However, together with the two SLSPs from Cryptomycota and Microsporidia, the 11 entomophthoralean proteases are a unique group of proteases exclusive to some of the early diverging fungal lineages.

Functional annotation indicates apparent protease activity based on sequence similarity, but function of the novel 11 SLSPs in group C is unknown. Eight of these SLSPs possess a signal peptide that suggest external secretion and thus indicative of a function on the immediate environment, whereas the remaining three might not be secreted or incomplete sequence models. Apart from the canonical protease S8 domain (PF00082), no other Pfam domains were found among this group C SLSPs. Searches against InterPro databases similarly did not reveal any other protein domains apart from the protease S8 SLSP domain (PRINTS: subtilisin serine protease family (S8) signature (PR00723), InterPro: peptidase S8, subtilisin-related (IPR015500), and ProSitePatterns: serine proteases, subtilase family (PS00138)).

Notably, none of these SLSPs contain the proteinase inhibitor i9 domain (PF05922) often found among the classical protease K-like SLSPs (Muszewska et al., 2011). However, this group C of SLSPs may potentially have evolved a different function than the “classical” fungal SLSPs in group A as evidenced by extensive diversification of the amino acids immediately surrounding the active site residues in the DHS triad (Figure 2). Out of the five Entomophthorales species analysed here, only three: *C. incongruus, E. muscae* and *P. formicae* contained members in the new group C SLSPs (Figure 1). Since these genes are missing in the genomes of *C. coronatus* and *C. thromboides*, the absence of particular SLSPs is unlikely to be due to sequencing or sampling artifacts. The unequal phylogenetic presence of the group C SLSPs could be indicative of specific functions related to niche adaptation. The two insect-pathogenic fungi specialized on house flies (*E. muscae*) and wood ants (*P. formicae*) contain five and two of the novel group C SLSPs, respectively (Figure 1), while no group C SLSP S8A-proteases were found in Basidiomycota or Ascomycota (Figure 3). However, the insect-pathogenic hypocrealean and nematode-trapping fungi within Ascomycota contain their own unique SLSP’s missing in Entomphthoromycotina (Figure 4). The soil saprobe and opportunistic human pathogen *C. incongruus* also contains four group C SLSPs suggesting these SLSPs are not exclusively related to host-specific evolution of the specialist insect-pathogenic entomophthoralean fungi.

**Figure 4.**
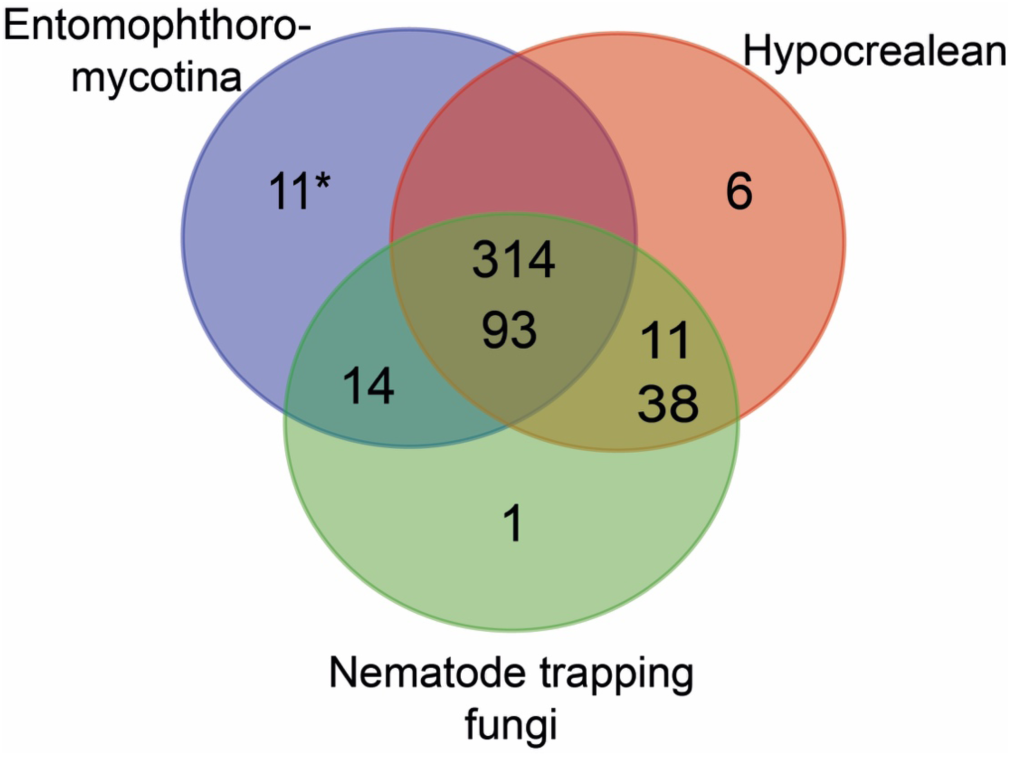
Venn diagram showing taxonomic distribution of subtilisin-like serine protease clusters of major insect and nematode-pathogenic fungal genera. Each two or three-digit number corresponds to number of S8A proteases in a specific cluster, which in two cases contain two clusters (314 and 93, and 11 and 38). The asterisk (*) marks the 11 members of the new bacillo-peptidase like cluster C found within the order Entomophthorales (Entomophthoromycotina). Entomophthoromycotina encompasses SLSP’s found in the genera: *Basidiobolus, Conidiobolus, Entomophthora,* and *Pandora,* Hypocrealean consists of SLSP’s from the genera: *Cordyceps, Metarhizium, Ophiocordyceps,* and Nematode trapping fungi is SLSP’s found in the genera: *Arthrobotrys* and *Monacrosporium.*

Further studies including genomic comparisons of the host-specific insect-pathogenic Entomophthorales will likely shed interesting new light on the gene content of these early diverging fungi (De Fine Licht et al., 2016). The presence of unusual genome organization, polyploidy and large genomes in many host-specific insect-pathogenic species within Entomophthorales has previously been a hindrance to genome sequencing (Gryganskyi and Muszewska, 2014). However, the present analysis exemplifies the many new proteins and enzymes that may be discovered as genomes begin to become available within Entomophthorales.

## Acknowledgements

The work conducted by the U.S. Department of Energy Joint Genome Institute is supported by the Office of Science of the U.S. Department of Energy under Contract No. DE-AC02-05CH11231. Work by JES and AG was partially supported by funding from the National Science Foundation (DEB-1441715) to JES. HHDFL was supported by the Villum Foundation (grant no. 10122).

